# Unique episymbiotic relationship between *Gracilibacteria* and *Zoogloea* in activated sludge flocs in a municipal wastewater treatment plant

**DOI:** 10.1101/2023.08.16.553637

**Authors:** Naoki Fujii, Kyohei Kuroda, Takashi Narihiro, Yoshiteru Aoi, Noriatsu Ozaki, Akiyoshi Ohashi, Tomonori Kindaichi

**Affiliations:** Department of Civil and Environmental Engineering, Graduate School of Advanced Science and Engineering, Hiroshima University, 1-4-1, Kagamiyama, Higashihiroshima, Hiroshima 739-8527, Japan; Bioproduction Research Institute, National Institute of Advanced Industrial Science and Technology (AIST), 2-17-2-1 Tsukisamu-Higashi, Toyohira-ku, Sapporo, Hokkaido 062-8517, Japan; Program of Biotechnology, Graduate School of Integrated Sciences for Life, Hiroshima University, 1-3-1, Kagamiyama, Higashihiroshima, Hiroshima 739-8530, Japan

**Author notes:** **Corresponding author:** Tomonori Kindaichi. **Competing Interests** The authors declare no competing interests.

## Abstract

Among the various bacteria present in activated sludge, uncultivated *Patescibacteria* (also known as the Candidate Phyla Radiation/CPR superphylum) are ubiquitous at the class or phylum level. *Patescibacteria* have a highly restricted metabolic capacity and are thought to be episymbiotic/endosymbiotic or predatory. However, only a limited number of *Patescibacteria* and their hosts have been identified. Therefore, many *Patescibacteria* have not been (co-)cultured and identified by fluorescence *in situ* hybridization (FISH) or electron microscopy. Little is known about the morphology, metabolic potential, and hosts of *Gracilibacteria* (formerly GN02 or BD1-5) which belong to *Patescibacteria*. In our previous study, we confirmed the presence of *Gracilibacteria* in activated sludge and successfully recovered its high-quality genome. In this study, we designed new probes to visualize members of *Gracilibacteria* in activated sludge and identified its host using FISH. The FISH observations revealed that *Gracilibacteria*, which formed loosely associated clusters, were located within dense clusters of *Zoogloea*, which were dominant in the activated sludge. The metagenome-assembled genomes (MAGs) of *Zoogloea* possessed genes related to extracellular polymeric substance (EPS) biosynthesis, floc formation, and nutrient removal, including a polyhydroxyalkanoate (PHA) accumulation pathway. The MAGs of *Gracilibacteria* possessed genes associated with type IV pili, competence protein EC (ComEC), and PHA degradation, which suggests that they have a *Zoogloea*-dependent lifestyle in activated sludge flocs. These findings clearly indicate a new symbiotic relationship between *Gracilibacteria* and *Zoogloea*, and to the best of our knowledge, this is the first study to show this interaction.

## Introduction

The activated sludge method is a biological treatment that has been widely used in wastewater treatment plants for over 100 years. This method removes organic matter and nutrients from influent wastewater through metabolic reactions of various microorganisms in activated sludge. However, the detailed taxonomic and physiological characteristics of the microbial communities within activated sludge, which are known for their complexity, are not well understood [1]. Among these diverse bacteria, several members of the bacterial phylum *Patescibacteria* (also known as the Candidate Phyla Radiation/CPR superphylum), a large and diverse bacterial group consisting of uncultivated bacteria, are universally present [1−4]. *Patescibacteria* are characterized by their limited metabolic potential, as inferred from metagenomic analyses [2, 5, 6]. Furthermore, microscopic observation using techniques such as fluorescence *in situ* hybridization (FISH) and transmission electron microscopy (TEM) have revealed that most *Patescibacteria* have small cell sizes. Recent studies have highlighted that *Patescibacteria* parasitize other bacterial or archaeal microorganisms, suggesting a lifestyle that may compensate for their metabolic deficiencies [7−12]. Within the *Patescibacteria* phylum, *Saccharimonadia, Paceibacteria*, and *Gracilibacteria* are commonly detected in the activated sludge flocs [1, 2, 5, 13], and information on *Gracilibacteria* has only recently been obtained by metagenomic analyses. To better understand microbial interactions in activated sludge flocs, *in situ* visualization using FISH or TEM within the flocs is very important because FISH probes specific for *Gracilibacteria* did not exist until recently. In this study, we designed new probes based on full-length 16S rRNA genes from previously obtained metagenome-assembled genomes (MAGs) related to *Gracilibacteria* from activated sludge flocs and visualized them using FISH. The FISH observations showed that *Gracilibacteria* surrounds *Zoogloea* cells in activated sludge flocs, indicating a symbiotic metabolic interaction. Therefore, the metabolic potentials of *Gracilibacteria* and *Zoogloea* in activated sludge flocs were predicted using the MAGs.

### FISH probes for *Gracilibacteria* and *Zoogloea*

We reconstructed one *Gracilibacteria* MAG and two *Zoogloea* MAGs from activated sludge flocs using a previous dataset (DRA013531) [2] (**Supplementary Table 1**). For the selection of the specific probes, 16S rRNA gene sequence was present in the MAG of *Gracilibacteria* but not in those of *Zoogloea*. Therefore, it was reconstructed using EMIRGE [14]. To detect *Gracilibacteria* in activated sludge flocs, we designed two new oligonucleotide probes (GRA665 and GRA686) (**Supplementary Table 2, Supplementary Fig. 1**). For the detection of *Zoogloea*, a previously reported probe, ZOO834 [15], was used because the probe sequence perfectly matched the reconstructed 16S rRNA gene of *Zoogloea*. The probe ZOO834 covered most *Zoogloea* species (89.7% of Zoogloea spp. based on SILVA v138.1 taxonomy), including the reconstructed full-length of 16S rRNA gene sequence of Zoogloea in this study (Supplementary Fig. 2).

### In situ detection and morphology of *Gracilibacteria* and *Zoogloea*

The cells hybridized with probes GRA665 and GRA686 showed loosely associated clusters of tiny cocci (< 0.5 μm in diameter) (**Fig. 1A**). These cells were also hybridized with bacteria-specific probes, EUBmix (**Fig. 1B**); therefore, the cells positive for probes GRA665 and GRA686 likely belonged to *Gracilibacteria. Gracilibacteria* formed loosely associated clusters within rod-shaped cells with dense clusters that hybridized with the EUBmix probe (**Fig. 1B**). The rod-shaped cells were likely *Zoogloea* spp. based on their unique finger-like shaped clusters. In fact, the rod-shaped cells were also hybridized with the previously designed *Zoogloea*-targeting probe, ZOO834 (**Fig. 1C**). Therefore, from the combination of GRA665/GRA686 and ZOO834 probes, we observed most *Gracilibacteria* cells in dense clusters of *Zoogloea* (**Fig. 1C and 1D**). Some members of *Paceibacteria* are known to localize to the cell poles of *Methanospirillum* [9], but the *Gracilibacteria* observed in this study did not adhere to specific sites on the *Zoogloea* cells (**Fig. 1D)**. *Zoogloea* is known to form dense cell aggregates, such as the formation of activated sludge flocs; interestingly, when *Gracilibacteria* formed dense clusters, the clusters of zoogloeal cells tended to decrease (**Fig. 1E−G**). Thus, *Gracilibacteria* may have a negative effect on *Zoogloea*. In addition, some of *Gracilibacteria* were present very close to *Zoogloea* cells that appeared to be undergoing cell division (**Fig. 1 H**). Previous studies on *Patescibacteria* [7−9] suggest that *Gracilibacteria* have symbiotic relationships with other bacteria as hosts. However, further studies are required to clarify *Gracilibacteria’s* effect on the growth condition of *Zoogloea*.

**Fig. 1.**
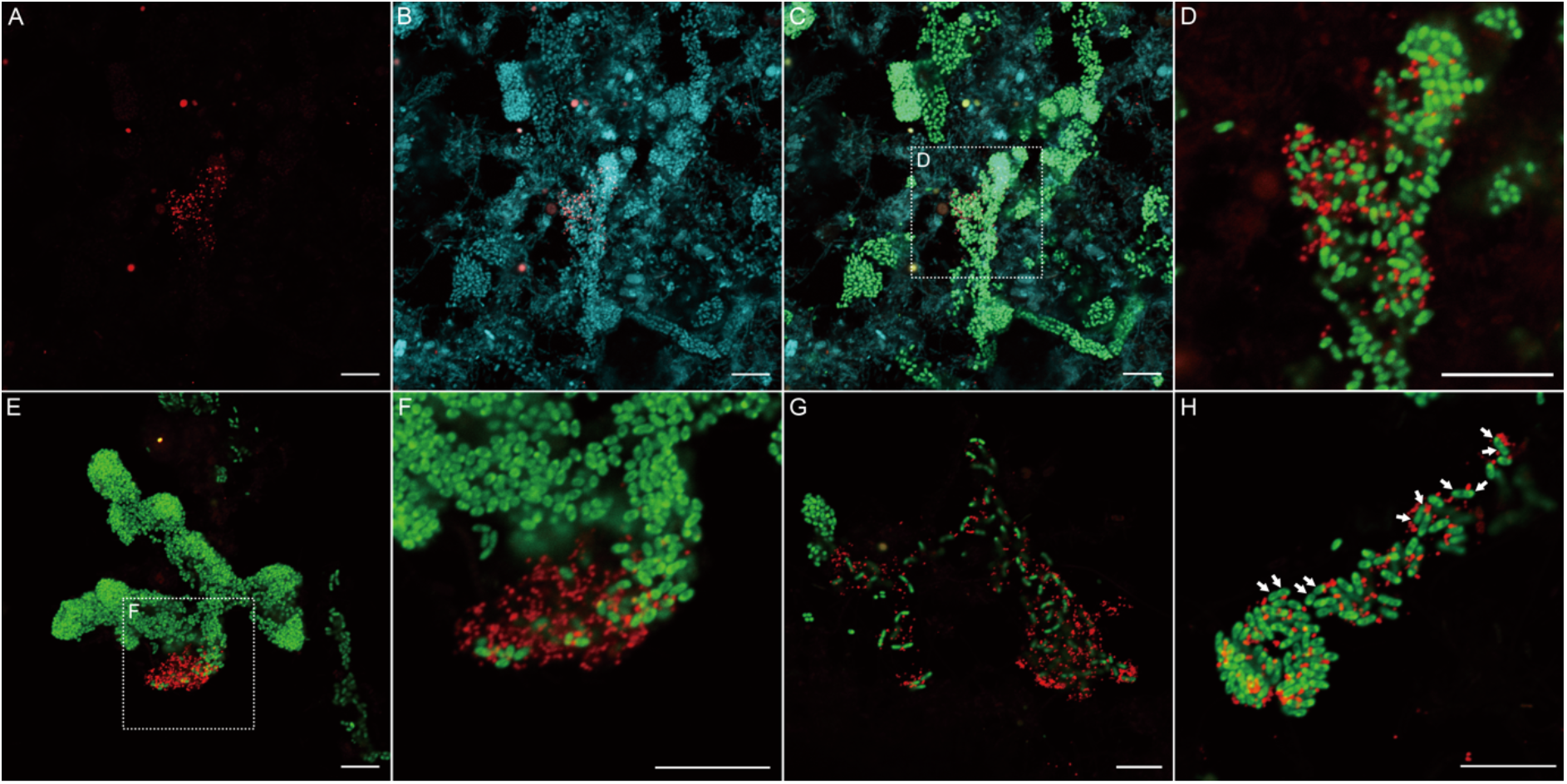
Fluorescence in situ hybridization (FISH) micrographs of *Gracilibacteria* and *Zoogloea* in activated sludge flocs. FISH was performed with three different fluorophores, the Alexa 488-labeled ZOO834 probe (green), Alexa 555-labeled GRA655 or GRA686 probe (red), Alexa 647-labeled EUBmix probes. (A-C) FISH micrographs of the same field of view; (A) GRA655 (red), ZOO834 (green), GRA655 (red), and EUBmix (blue), (B) GRA655 (red), and EUBmix (blue), (C) ZOO834 (green), GRA655 (red), and EUBmix (blue), (D) ZOO834 (green) and GRA655 (red). (E and F) ZOO834 (green) and GRA686 (red). (G and H) ZOO834 (green) and GRA655 (red). White arrows in (H) indicate zoogloeal cells undergoing cell division. Scale bars = 10 μm.

### The relationship between *Gracilibacteria* and *Zoogloea* predicted from the genomic traits

*Zoogloea* is known to be involved in floc formation in the activated sludge by secretion extracellular polymeric substance (EPS). In the two reconstructed *Zoogloea* MAGs (**Fig. 2A**), we identified homologs of clusters related to EPS biosynthesis and export and floc formation [16, 17] (**Fig. 2B**). We identified most of the critical genes (shown as pink in **Fig. 2B**) for each function in the *Zoogloea* MAGs and other functional genes for polyhydroxyalkanoate (PHA) accumulation, denitrification, sulfur metabolism, and glycogen accumulation, suggesting that *Zoogloea* may be involved in nutrient removal and floc formation within activated sludge (**Supplementary Information**).

**Fig. 2.**
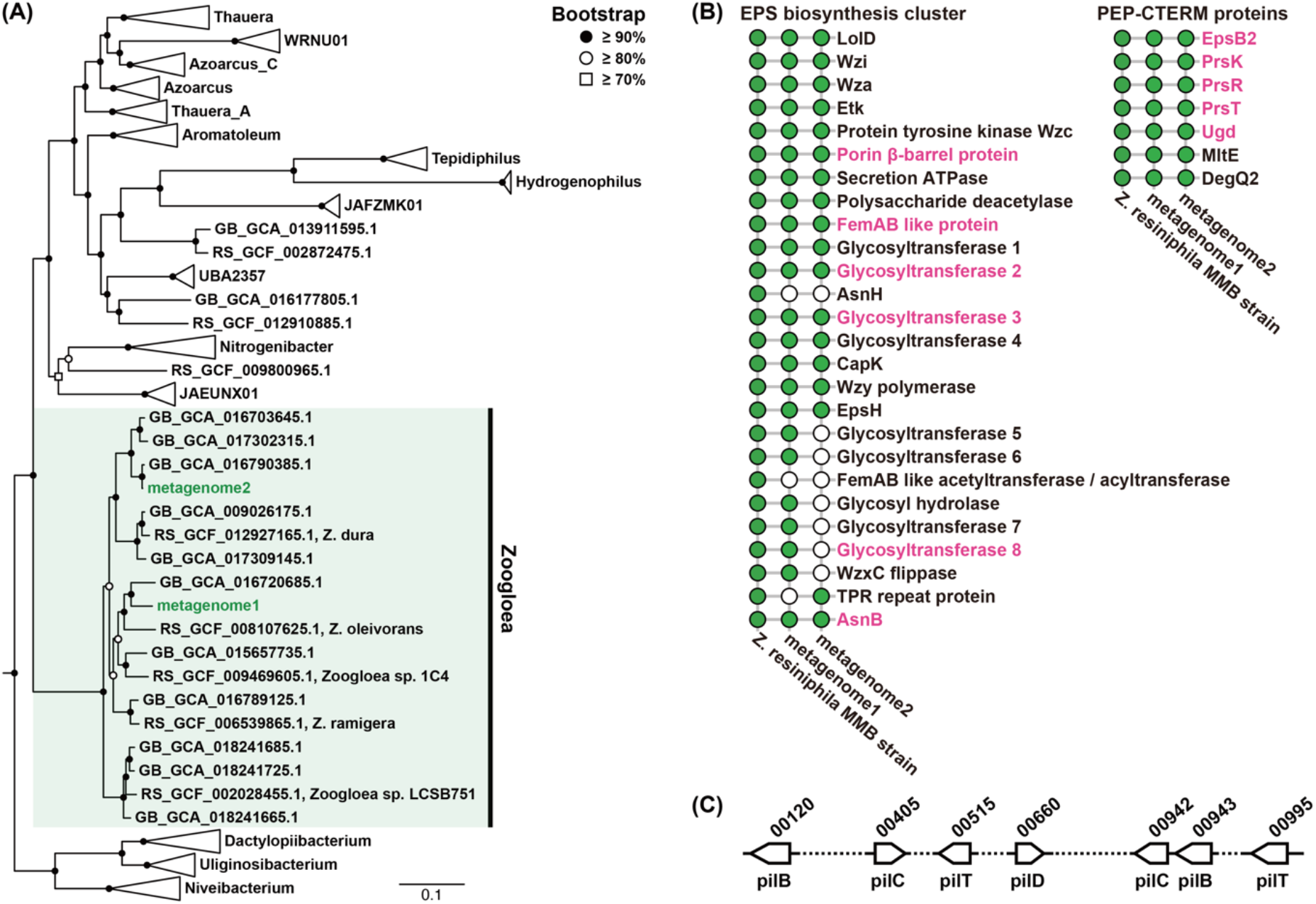
Phylogenetic and important genes analysis. (A) Phylogenetic tree of the family *Rhodocyclaceae* based on concatenated phylogenetic marker genes of GTDB-Tk v2.2.6 (R207). The phylogenetic position of the metagenomic bins (green). (B) The presence of the extracellular polymeric substance (EPS) biosynthesis cluster genes [16] and phosphoenolpyruvate (PEP)-CTERM proteins [17] in the metagenomic bins. Proteins (in pink) indicate those critical in each cluster. Green circles indicate identified genes and white circles indicate genes could not be identified in this study. (C) The location of genes associated with type IV pili and locus tags.

In *Gracilibacteria*, we identified genes involved in type IV pili (pilB, -C, -D, and -T), which are common in many *Patescibacteria* (**Fig. 2C**). Type IV pili have recently been shown to be involved in adhesion to host bacteria and host-specific recognition [12] and have been implicated in DNA uptake through conjugation with the competence protein EC (ComEC) gene [18]. Since Grac*ilibacteria* also possess the ComEC gene, it may be involved in these roles. In addition, DNA, polysaccharides, and other macromolecules are found in the EPS fraction of activated sludge [19]. Because *Gracilibacteria* also possess PHA depolymerase they could possibily form a symbiotic relationship with *Zoogloea*. Since *Zoogloea* may also be involved in PHA accumulation, they can create an environment favorable for survival of *Gracilibacteria*. Another unique feature of *Gracilibacteria* is that they possess pyruvate kinase in their glycolysis pathway. No membrane transporters such as phosphoenolpyruvate (PEP) were observed. Thus, whether the *Gracilibacteria* incorporates PEP from outside the cell or internally synthesizes it remains to be clarified. These findings strongly suggest a novel episymbiotic lifestyle of *Patescibacteria* on the *Zoogloea*-formed flocs, and this episymbiotic relationship must be investigated in more detail for the control and design of microbial ecology in activated sludge.

## Supporting information

Supplementary data sheet

## Acknowledgements

This work was supported by JSPS KAKENHI [Grant No. JP16H04833 and JP20H02287].

## Data Availability Statement

The sequence data of the partial 16S rRNA gene sequence were deposited in the GenBank/EMBL/DDBJ databases under the accession number DRA013509. Metagenomic sequence data were deposited in the DDBJ database under the DDBJ/EMBL/GenBank accession number DRA013531.

