## Supplementary Information for "Unique episymbiotic relationship between *Gracilibacteria* and *Zoogloea* in activated sludge flocs in a municipal wastewater treatment plant"

### Materials and Methods

#### Sample collection

As previously described [1], activated sludge samples were collected from aeration tanks in a wastewater treatment plant in Higashihiroshima City in February 2019 (designated as AS201902) and April 2020 (designated as AS202004). The AS202004 sample was anaerobically incubated for 3 d to change the relative abundance of *Patescibacteria* and was then designated as AA202004. Fresh and incubated sludge samples were stored at –18°C for further analyses.

#### 16S rRNA gene amplicon sequences and metagenomic analysis

16S rRNA gene amplicon sequencing results were obtained from a previous study [1] using QIIME2 v2021.11 [2] and the SILVA v138.1 database [3]. Full-length 16S rRNA gene sequences of *Zoogloea* were reconstructed using EMIRGE software (–151-i 350-s 50-phred33). [4] The 16S rRNA gene sequence was compared with the partial length of amplicon sequence variants (ASVs) of *Zoogloea* (**Supplementary Table 3**). The partial sequence of the most abundant zoogloeal ASV corresponded to the reconstructed 16S rRNA gene with 100% identity (**Supplementary Fig. 2**). Metagenome-assembled genomes (MAGs) of *Gracilibacteria* and *Zoogloea* (DRA013531) were obtained from a previous study [1]. Phylogenetic classification of *Zoogloea* was evaluated using GTDB-Tk v2.2.6 [5]. Genomes were annotated using a combination of Prokka v1.13, [6] GhostKOALA [7], and manual annotation. The genome tree was constructed by using a combination of GTDB-Tk and IQ-TREE v2.2.2.3. Conserved marker genes were identified using "gtdbtk identify" as the default parameter, and a phylogenetic filter (--taxa_filter f__Rhodocyclaceae) was applied with "gtdbtk align" and aligned to the reference genome using "gtdbtk align". Phylogenetic trees were constructed using an auto-optimized surrogate model (Q.insect+F+R10) in IQ-TREE (B 1000) [8]. The two reconstructed *Zoogloea* MAGs did not contain the 16S rRNA gene; thus, they were classified into the genus *Zoogloea* in a phylogenetic tree based on concatenated phylogenetic marker genes (**Fig. 2A**). Therefore, we speculated that both the reconstructed full-length 16S rRNA gene and genomes were derived from dominant *Zoogloea* in activated sludge flocs. Zoogloeal MAGs were assigned to the reference genome of *Z. resiniphila* MMB strain (KX259245) [9] using BLASTp v2.6.0 [10] for manual annotation.

**Probe design and fluorescence *in situ* hybridization**

Probes specific for Gracilibacteria were designed using ARB software v7.0 [11]. The specificity and coverage of the newly designed GRA665 and GRA686 are shown in **Supplementary Fig. 1**. The optimal formamide concentrations of GRA665 and GRA686 were 30% each, which was predicted using mathFISH [12] (**Supplementary Figs. 3 and 4, and Supplementary Table 2**). GRA665 is a highly specific probe, and can detect only one sequence with the targeted sequence in the SILVA v138.1 database. In contrast, GRA686 can detect clusters in the SILVA v138.1 database, including closely related sequences. GRA686 was predicted to match out-of-class sequences with a single nucleotide mismatch; however, mathFISH [12] predicted that GRA686 did not hybridize to off-target bacteria under optimal formamide concentrations [13]. (**Supplementary Fig. 4**). Furthermore, adding a competitor probe almost completely prevents false positive hybridization; therefore, using a competitor probe is recommended (**Supplementary Fig. 5**). Sample fixation and FISH were performed as previously described [14]; the AS202004 sample was used for FISH. The probes used in this study are listed in **Supplementary Table 2**. Probes were labeled at the 5' end with either Alexa488, Alexa555, or Alexa647. Simultaneous hybridization was performed using the procedure described below. Hybridization with the probe requiring higher stringency was performed first, and then hybridization with the probe requiring lower stringency was performed [15]. The FISH observation was performed with a LSM700 confocal laser-scanning microscope equipped with lasers with excitation wavelengths of 488, 555, and 639 nm.

**Supplementary Figures**

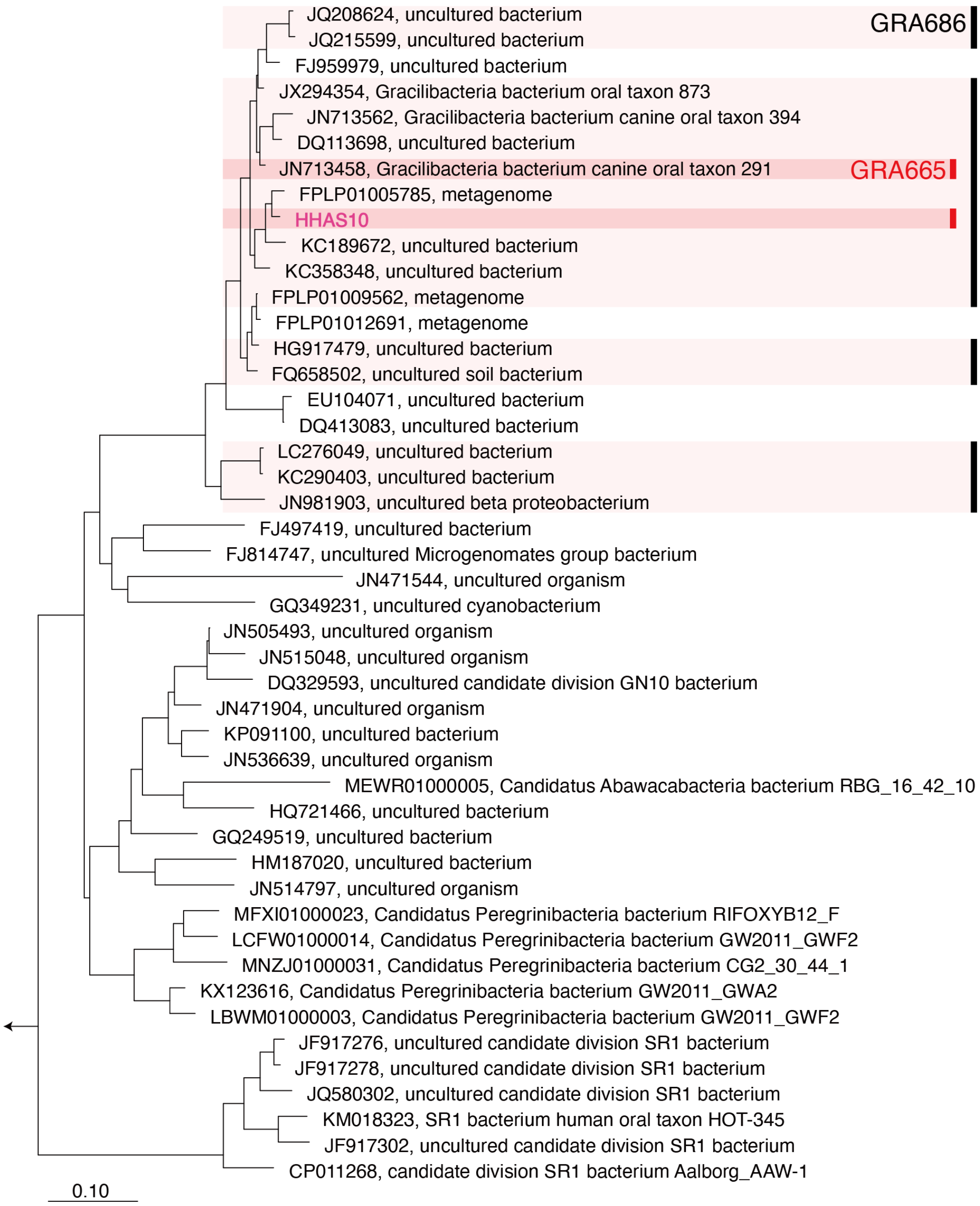


**Supplementary Fig. 1**

Maximum likelihood phylogenetic tree of *Gracilibacteria*-related 16S rRNA genes. Sequences named HHAS10 (in pink) indicate 16S rRNA genes obtained from the previous metagenomic analysis [1]. The vertical, colored lines for the probes GRA665 and GRA686 indicate the sequences possibly hybridized with the probe with zero mismatches. 1000-replicate bootstraps were used.

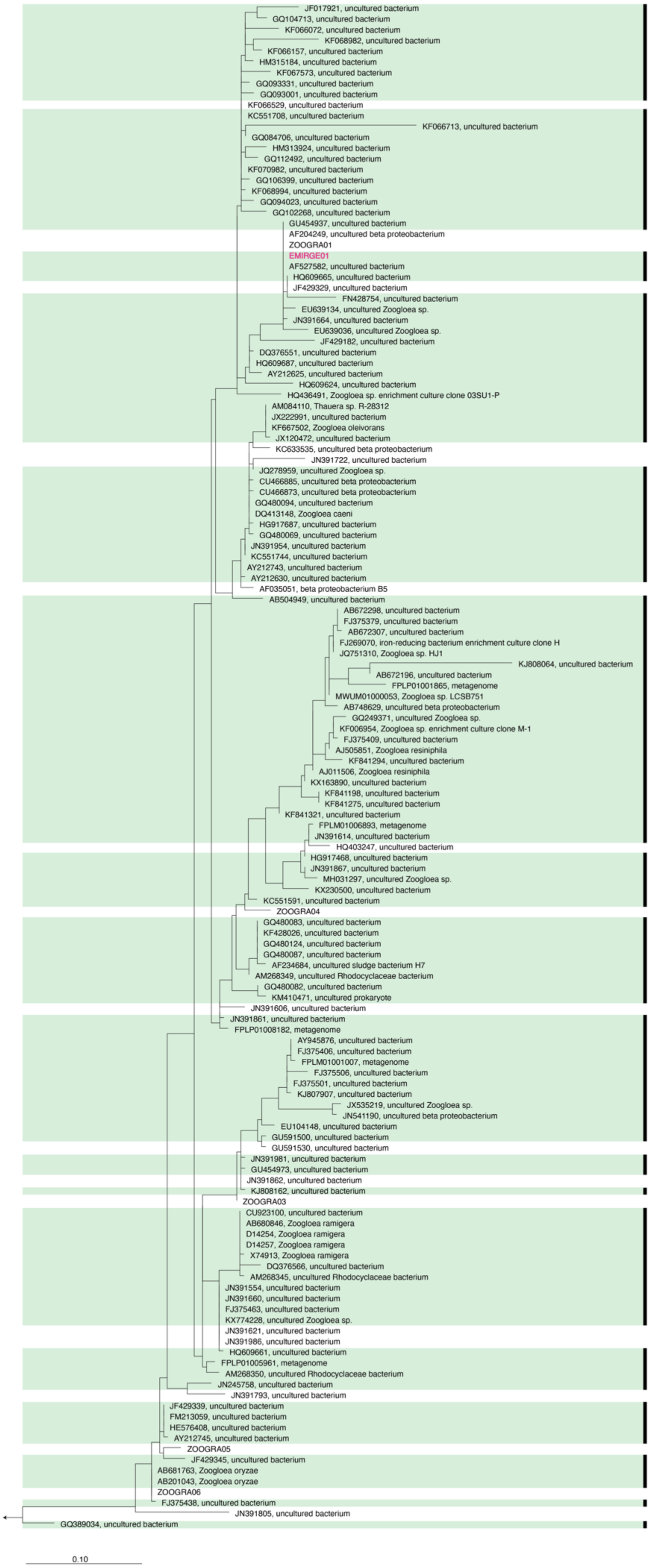
JF017921, uncultured bacterium
GQ104713, uncultured bacterium
KF066072, uncultured bacterium
KF068982, uncultured bacterium
KF066157, uncultured bacterium
HM315184, uncultured bacterium
KF067573, uncultured bacterium
GQ093331, uncultured bacterium
GQ093001, uncultured bacterium
KF066529, uncultured bacterium
KC551708, uncultured bacterium
KF066713, uncultured bacterium
GQ084706, uncultured bacterium
HM313924, uncultured bacterium
GQ112492, uncultured bacterium
KF070982, uncultured bacterium
GQ106399, uncultured bacterium
KF068994, uncultured bacterium
GQ094023, uncultured bacterium
GQ102268, uncultured bacterium
GU454937, uncultured bacterium
AF204249, uncultured beta proteobacterium
ZOOGRA01
EMIRGE01
AF527582, uncultured bacterium
HQ609665, uncultured bacterium
JF429329, uncultured bacterium
FN428754, uncultured bacterium
EU639134, uncultured Zoogloea sp.
JN391664, uncultured bacterium
EU639036, uncultured Zoogloea sp.
JF429182, uncultured bacterium
DQ376551, uncultured bacterium
HQ609687, uncultured bacterium
AY212625, uncultured bacterium
HQ609624, uncultured bacterium
HQ436491, Zoogloea sp. enrichment culture clone 03SU1-P
AM084110, Thauera sp. R-28312
JX222991, uncultured bacterium
KF667502, Zoogloea oleivorans
JX120472, uncultured bacterium
KC633535, uncultured beta proteobacterium
JN391722, uncultured bacterium
JQ278959, uncultured Zoogloea sp.
CU466885, uncultured beta proteobacterium
CU466873, uncultured beta proteobacterium
GQ480094, uncultured bacterium
DQ413148, Zoogloea caeni
HG917687, uncultured bacterium
GQ480069, uncultured bacterium
JN391954, uncultured bacterium
KC551744, uncultured bacterium
AY212743, uncultured bacterium
AY212630, uncultured bacterium
AF035051, beta proteobacterium B5
AB504949, uncultured bacterium
AB672298, uncultured bacterium
FJ375379, uncultured bacterium
AB672307, uncultured bacterium
FJ269070, iron-reducing bacterium enrichment culture clone H
JQ751310, Zoogloea sp. HJ1
KJ808064, uncultured bacterium
AB672196, uncultured bacterium
FPLP01001865, metagenome
MWUM01000053, Zoogloea sp. LCSB751
AB748629, uncultured beta proteobacterium
GQ249371, uncultured Zoogloea sp.
KF006954, Zoogloea sp. enrichment culture clone M-1
FJ375409, uncultured bacterium
AJ505851, Zoogloea resiniphila
KF841294, uncultured bacterium
AJ011506, Zoogloea resiniphila
KX163890, uncultured bacterium
KF841198, uncultured bacterium
KF841275, uncultured bacterium
KF841321, uncultured bacterium
FPLM01006893, metagenome
JN391614, uncultured bacterium
HQ403247, uncultured bacterium
HG917468, uncultured bacterium
JN391867, uncultured bacterium
MH031297, uncultured Zoogloea sp.
KX230500, uncultured bacterium
KC551591, uncultured bacterium
ZOOGRA04
GQ480083, uncultured bacterium
KF428026, uncultured bacterium
GQ480124, uncultured bacterium
GQ480087, uncultured bacterium
AF234684, uncultured sludge bacterium H7
AM268349, uncultured Rhodocyclaceae bacterium
GQ480082, uncultured bacterium
KM410471, uncultured prokaryote
JN391606, uncultured bacterium
JN391861, uncultured bacterium
FPLP01008182, metagenome
AY945876, uncultured bacterium
FJ375406, uncultured bacterium
FPLM01001007, metagenome
FJ375506, uncultured bacterium
FJ375501, uncultured bacterium
KJ807907, uncultured bacterium
JX535219, uncultured Zoogloea sp.
JN541190, uncultured beta proteobacterium
EU104148, uncultured bacterium
GU591500, uncultured bacterium
GU591530, uncultured bacterium
JN391981, uncultured bacterium
GU454973, uncultured bacterium
JN391862, uncultured bacterium
KJ808162, uncultured bacterium
ZOOGRA03
CU923100, uncultured bacterium
AB680846, Zoogloea ramigera
D14254, Zoogloea ramigera
D14257, Zoogloea ramigera
X74913, Zoogloea ramigera
DQ376566, uncultured bacterium
AM268345, uncultured Rhodocyclaceae bacterium
JN391554, uncultured bacterium
JN391660, uncultured bacterium
FJ375463, uncultured bacterium
KX774228, uncultured Zoogloea sp.
JN391621, uncultured bacterium
JN391986, uncultured bacterium
HQ609661, uncultured bacterium
FPLP01005961, metagenome
AM268350, uncultured Rhodocyclaceae bacterium
JN245758, uncultured bacterium
JN391793, uncultured bacterium
JF429339, uncultured bacterium
FM213059, uncultured bacterium
HE576408, uncultured bacterium
AY212745, uncultured bacterium
ZOOGRA05
JF429345, uncultured bacterium
AB681763, Zoogloea oryzae
AB201043, Zoogloea oryzae
ZOOGRA06
FJ375438, uncultured bacterium
JN391805, uncultured bacterium
GQ389034, uncultured bacterium
0.10

**Supplementary Fig. 2**

Maximum likelihood phylogenetic tree of *Zoogloea*-related 16S rRNA genes. Sequences named EMIRGE01 (in pink) indicate 16S rRNA gene reconstructed with EMIRGE software. Sequences named ZOOGRA* are ASVs obtained from 16S rRNA amplicon sequence analysis. The vertical, colored lines for each probe indicate the OTUs possibly hybridized with the probe ZOO834 with zero mismatch. 1000-replicate bootstraps were used.

***RESULTS : General Analysis***

```
PROBE ALIGNMENT :

...........................
PROBE--3'CCCGATCGTCTTGCCATT5'
.........||||||||||||||||||
TARGET-5'GGGCTAGCAGAACGGTAA3'
...........................
```

| | |
|---|---|
| $\Delta G^o_1$ | -21.9 kcal/mol |
| $\Delta G^o_2$ | 1.8 kcal/mol |
| $\Delta G^o_3$ | -7.5 kcal/mol |
| $\Delta G^o_{overall}$ | -14.4 kcal/mol |
| $[FA]_m$* | 31.2 % |
| Hybridization Efficiency** | 0.9994 |

* Melting formamide concentration
** At 0% formamide

### HYBRIDIZATION EFFICIENCY CURVE

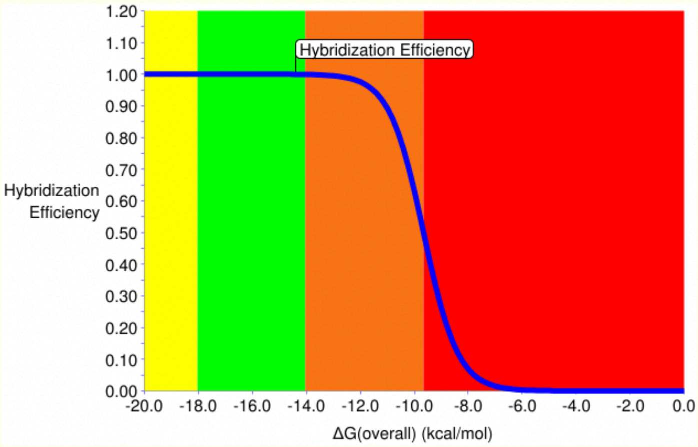


Hybridization Efficiency Meter !!

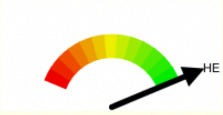


### FORMAMIDE CURVE

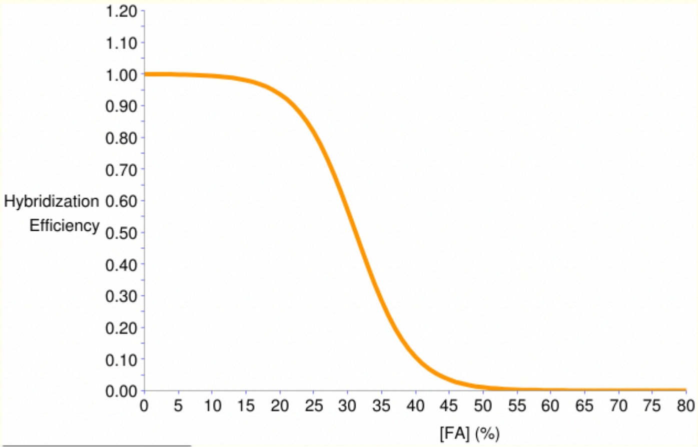


**Supplementary Fig. 3**

mathFISH curve for the GRA665 probe. The target organism is HHAS10.

#### *RESULTS : Mismatch Analysis*

```
PROBE ALIGNMENT WITH TARGET ORGANISM:
...........................
PROBE--3'ATCCCCACGTTAGGCAAC5'
.........||||||||||||||||||
TARGET-5'TAGGGGTGCAATCCGTTG3'
...........................

PROBE ALIGNMENT WITH NON TARGET ORGANISM:
.................G.........
PROBE--3'ATCCCCAC.TTAGGCAAC5'
.........||||||||.|||||||||
TARGET-5'TAGGGGTG.AATCCGTTG3'
.................A.........
```

| | TARGET ORGANISM | NON-TARGET ORGANISM | ΔValue(?) |
|---|---|---|---|
| $\Delta G^o_1$ | -20.0 kcal/mol | -16.2 kcal/mol | 3.80 kcal/mol |
| $\Delta G^o_2$ | 0.8 kcal/mol | 0.8 kcal/mol | NA |
| $\Delta G^o_3$ | -4.9 kcal/mol | -6.4 kcal/mol | -1.50 kcal/mol |
| $\Delta G^o_{overall}$ | -15.0 kcal/mol | -9.7 kcal/mol | 5.30 kcal/mol |
| $[FA]_m$* | 30.0 % | 0.1 % | -29.90 % |
| Hybridization Efficiency** | 0.9998 | 0.4998 | -0.50 |

* Melting formamide concentration
** At 0% formamide

### FORMAMIDE CURVE

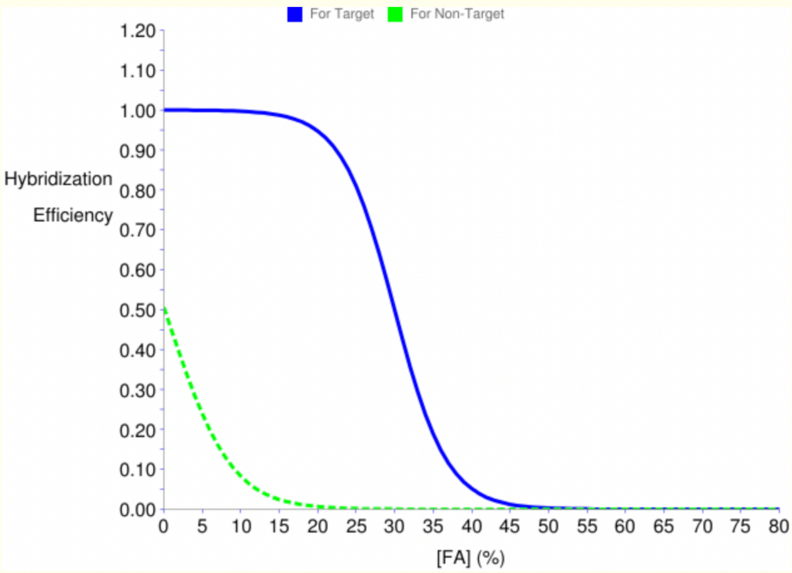


**Supplementary Fig. 4**

mathFISH curve for the GRA686 probe. The target organism is HHAS10, and the non-target organism is AY168743.

### *RESULTS : Competitor Analysis*

*****
**Check: Probe is a perfect match to target, and has 1 mismatches to non-target.**
**Competitor is a perfect match to non-target and has 1 mismatches to target**

| | Probe - Target | Probe - Non Target | Competitor - Target | Competitor - Non Target |
|---|---|---|---|---|
| **$\Delta G^o_1$ kcal/mol** | -20.02 | -16.18 | -15.75 | -18.81 |
| **$\Delta G^o_2$ kcal/mol** | 0.8 | 0.8 | 1.15 | 1.15 |
| **$\Delta G^o_3$ kcal/mol** | -4.86 | -6.36 | -4.86 | -6.36 |
| **$\Delta G^o_{overall}$ kcal/mol** | -15.0 | -9.65 | -10.8 | -12.35 |
| **Hybridization Efficiency*** | 1.0 | 0.5 | 0.86 | 0.99 |

***At 0% formamide**

### *FORMAMIDE CURVE*

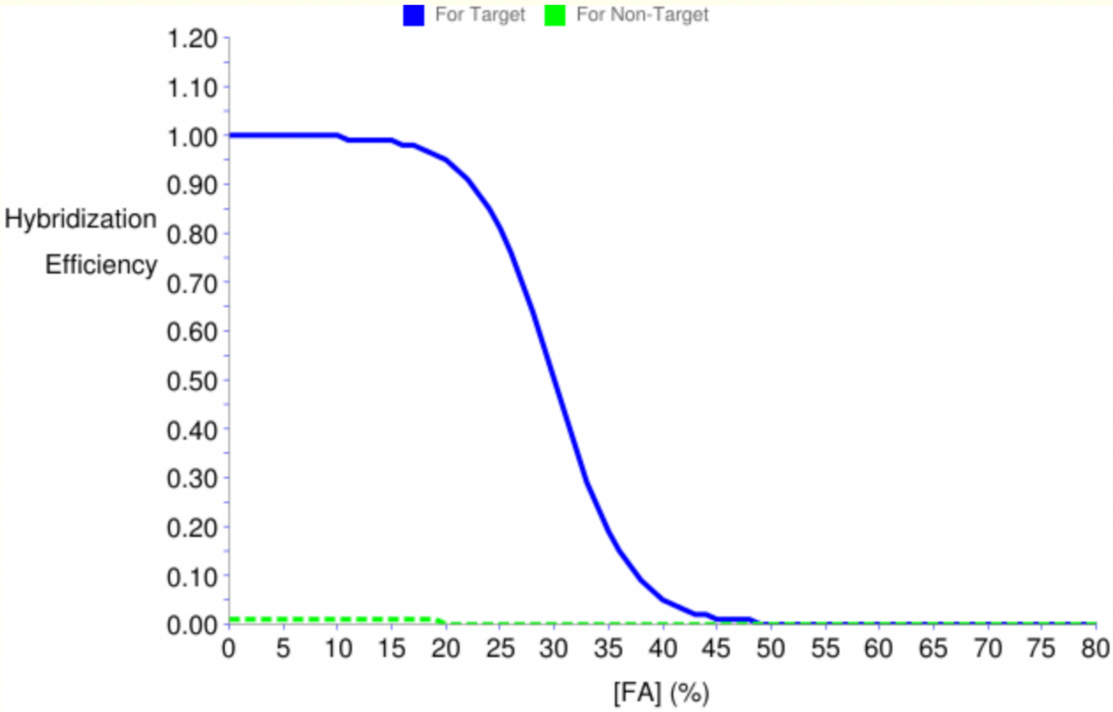


**Supplementary Fig. 5**

mathFISH curve for the GRA686 probe with a competitor probe. The target organism is HHAS10, and the non-target organism is AY168743.

### Supplementary Tables

**Supplementary Table 1** Basic information on the reconstructed *Gracilibacteria* and *Zoogloea* genomes.

| | **Taxonomy** | **Abundance** | | | **Quality Estimation** | | | **Bin Size (Mbp)** |
|---|---|---|---|---|---|---|---|---|
| | | **AS201902** | **AS202004** | **AA202004** | **Completeness** | **Contamination** | **Strain heterogeneity** | |
| HHAS10 | *Gracilibacteria* | 0.02 | 0.22 | 0.19 | 97.67* | 0* | 0* | 1.3 |
| metagenome1 | *Zoogloea* | 0.39 | 1.94 | 0.64 | 92.48 | 5.03 | 28.57 | 4.5 |
| metagenome2 | *Zoogloea* | 0.02 | 0.39 | 0.31 | 72.41 | 3.45 | 0 | 3.8 |

*Calculated using the CPR marker set.

**Supplementary Table 2**. The names, targets, sequences, formamide concentrations, and references of oligonucleotide probes used in this study.

| Probe name | Target | Sequence (5'→3') | FA (%) | Reference |
|---|---|---|---|---|
| GRA665 | Some members of *Gracilibacteria* | TTA CCG TTC TGC TAG CCC | 30 | This study |
| GRA686 | Some members of *Gracilibacteria* | CAA CGG ATT GCA CCC CTA | 30 | This study |
| CompGRA686 | Competitor for GRA686 | CAA CGG ATT TCA CCC CTA | - | This study |
| ZOO834 | Most member of *Zoogloea* | CTC AAT GAG TCT CCT CAC CG | 50 | 16 |
| EUB338 | Most *Bacteria* | GCT GCC TCC CGT AGG AGT | 0-50 | 17 |
| EUB338II | *Planctomycetales* | GCA GCC ACC CGT AGG TGT | 0-50 | 18 |
| EUB338III | *Verrucomicrobiales* | GCT GCC ACC CGT AGG TGT | 0-50 | 18 |
| EUB338IV | Bacteria lineages not covered by probes EUB338, EUB338II, and EUB338III | GCA GCC TCC CGT AGG AGT | 0-50 | 19 |

**Supplementary Table 3** Table of ASVs of *Zoogloea* obtained using QIIME2. The relative abundance of each sample is shown.

| | AS201902 (%) | AS202004 (%) | AA202004 (%) |
|---|---|---|---|
| ZOOGRA01 | 0.06 | 3.82 | 1.58 |
| ZOOGRA03 | 0 | 0.45 | 0.55 |
| ZOOGRA04 | 0 | 0.30 | 0.18 |
| ZOOGRA05 | 0 | 0.05 | 0.19 |
| ZOOGRA06 | 0.02 | 0.13 | 0 |
